# Computational Synthetic Inner Membrane Reveals Cardiolipin-Leak Control of ATP Output

**DOI:** 10.64898/2026.02.25.708092

**Authors:** Mark I.R. Petalcorin

## Abstract

The **inner mitochondrial membrane (IMM)** is a densely packed bioenergetic surface where **electron transfer, proton pumping**, and **ATP synthesis** are tightly coupled, and where performance is shaped by **membrane leak**, lipid composition, and higher-order organization. Although experimental reconstruction of oxidative phosphorylation in **proteoliposomes** has advanced, systematic exploration of design tradeoffs remains challenging because key parameters covary across preparations. Here we present a fully reproducible computational **synthetic IMM (syn-IMM)** framework that models coupled membrane energization and ATP production, then uses structured perturbations, parameter sweeps, and sensitivity analyses to identify dominant control variables. In a Δψ-proxy simulator, we show that ATP output is maximized within a narrow **cardiolipin performance window** (peak at cardiolipin fraction 0.18 in our benchmarked parameterization), while increasing **leak** globally suppresses performance, producing a cardiolipin × leak landscape in which coupling integrity is a primary gate. Perturbation experiments separate mechanistic regimes, reduced ATP synthase capacity yields an “energized but unproductive” state with preserved Δψ but depressed ATP flux, whereas increased leak reduces usable output despite relatively preserved energization. Monte Carlo sensitivity analysis ranks Δψ and respiratory capacity as strongest correlates of ATP output, with ATP synthase capacity contributing positively and leak contributing negatively. A multi-state extension introduces explicit **Δψ-ΔpH partitioning, finite CoQ redox pool dynamics**, and cardiolipin-dependent **supercomplex fraction S(t)**, enabling diagnosis of organization kinetics and driving-force partition effects. Together, this syn-IMM platform provides an interpretable bridge between component-level reconstruction and system-level design, offering quantitative acceptance tests and design rules for programmable bioenergetic membranes.

## Introduction

The **inner mitochondrial membrane (IMM)** is the cell’s most specialized **energy-converting surface**, an ultrastructured interface where **electron transfer** is converted into a stored electrochemical driving force and then into **ATP**, the dominant currency for cellular work. Unlike membranes whose primary roles are signaling or compartmentalization, the IMM is optimized for **high-flux bioenergetics**, packing an extraordinary density of macromolecular machines into a tightly regulated lipid environment (Brand & Nicholls, 2011). This density is not merely a descriptive feature, it is a functional requirement, because the IMM must sustain rapid turnover of redox reactions, maintain a stable driving force, and coordinate energy conversion with metabolite exchange. Function therefore emerges from **system-level coupling**, not from any single complex operating in isolation.

At the heart of IMM energy conversion is **oxidative phosphorylation (OXPHOS)**, which couples the **electron transport chain** to **proton pumping** and ultimately to ATP formation by **F**o**F**1**-ATP synthase**. Electron transfer through membrane complexes drives vectorial proton translocation, creating the **proton motive force (Δp)**. In bioenergetics, Δp is commonly expressed as an electrical equivalent, **Δp ≈ Δψ + 60·ΔpH (mV)**, where **Δψ** is the **membrane potential** and **ΔpH** is the transmembrane pH gradient (Perry et al., 2011). Although Δψ and ΔpH are thermodynamically equivalent contributors to Δp, they are not kinetically interchangeable in real systems. They can change on different timescales and respond differently to changes in buffering, ion transport, and membrane conductance, which means that relying on Δψ alone can obscure mechanistic regimes, especially in synthetic or reconstituted systems where ionic composition and buffering are deliberately manipulated (Perry et al., 2011). This is a central motivation for treating **Δψ** and **ΔpH** as separable contributors when building a synthetic IMM design logic.

A second defining principle is that the IMM is a biologically encoded hybrid of nuclear and mitochondrial genetic programs. The original sequencing and organization of the human mitochondrial genome established that mtDNA encodes a compact set of **13 OXPHOS polypeptides**, along with rRNAs and tRNAs required for mitochondrial translation (Anderson et al., 1981). Those 13 proteins are not peripheral accessories, they are the most hydrophobic, membrane-embedded cores of complexes I, III, IV, and V, positioned directly within the machinery that couples redox chemistry to proton translocation. This architecture is crucial for synthetic reconstruction because it highlights why mitochondrial bioenergetics is difficult to “rebuild” from scratch. The deeply hydrophobic, membrane-integrated subunits require coordinated insertion and assembly within a specialized lipid environment. As a result, most practical syn-IMM strategies, whether experimental or computational, begin by treating respiratory complexes as assembled functional modules rather than attempting de novo expression and insertion of mtDNA-encoded cores.

The IMM’s lipid environment is itself a determinant of function. Among IMM lipids, **cardiolipin** is distinctive in both enrichment and functional coupling to respiratory machinery. Cardiolipin is not simply a bulk component that fills space, it behaves as a stabilizing cofactor for OXPHOS assemblies and influences supramolecular organization. A foundational biochemical result demonstrated that cardiolipin is required for robust formation of **respiratory supercomplexes**, effectively “gluing” the respiratory chain into higher-order assemblies (Zhang et al., 2002). Subsequent evidence showed cardiolipin stabilizes respiratory chain supercomplexes in mitochondria, reinforcing the concept that lipid composition shapes functional organization rather than passively surrounding it (Pfeiffer et al., 2003). Cardiolipin-dependent organization has also been supported by reconstitution work demonstrating that respiratory supercomplex-like assemblies can be rebuilt from purified components in a cardiolipin-dependent manner (Bazán et al., 2013), and the broader functional implications of cardiolipin for mitochondrial supercomplex formation have been synthesized in later analyses (Mileykovskaya & Dowhan, 2014). Together, these studies motivate treating cardiolipin as a **tunable control knob** that can shift a membrane between poorly organized, weakly coupled states and more efficient, stable states.

IMM function is also inseparable from its architecture. The IMM forms cristae that expand surface area and reshape reaction-diffusion geometry, creating microenvironments where coupling can be reinforced or weakened depending on organization. A striking example of architecture arising from core machinery is that **ATP synthase dimers** can self-assemble into rows and induce membrane curvature, providing a direct causal link from an energy-converting enzyme to membrane shape (Blum et al., 2019). This observation matters for synthetic design because it implies that organization is not only a biochemical state but also a physical one. If organization changes membrane curvature and local packing density, it can change effective kinetics, diffusion constraints, and the stability of Δp. A syn-IMM framework that aims to explain performance must therefore consider organization as more than a static label.

A further layer often hidden in simplified depictions is the finite nature of mobile redox carriers, especially **coenzyme Q (CoQ)**. CoQ is a lipid-phase shuttle that connects multiple electron entry points to downstream oxidation, and in any reconstituted membrane it exists as a **finite pool** rather than an infinite bath. Sensitive quantitative work has demonstrated assays for CoQ pool redox state and emphasized its biological interpretability (Burger et al., 2020). In a finite-pool regime, CoQ can saturate and buffer flux, or become limiting and induce bottlenecks that alter how quickly a system reaches steady state and how it responds to perturbations. These considerations become especially important in proteoliposome reconstruction, where CoQ content can be precisely adjusted.

These principles converge on a timely opportunity. The ability to reconstruct mammalian respiratory function in defined membranes has advanced to the point where multi-component systems can be assembled and interrogated systematically. In particular, de- and reconstruction of the mammalian respiratory chain provides an experimental anchor that makes a “synthetic IMM” concept operational rather than speculative (Rimle et al., 2025). However, systematic exploration remains challenging because many experimental variables covary, including membrane leakiness, protein ratio and orientation, lipid composition, and carrier pool state. A computational syn-IMM framework can therefore serve as a bridge between biochemical reconstruction and systems-level interpretation, providing controlled perturbations, sensitivity analyses, and mechanistic diagnostics that guide experimental design.

In this work, we expand the synthetic IMM problem from a parts list to a **systems engineering** question. We focus on three mechanistic levers that are strongly supported by mitochondrial biology and directly relevant to reconstitution. First, the **partitioning of Δp** into **Δψ** and **ΔpH**, which shapes interpretation of coupling and output (Perry et al., 2011). Second, the finite and saturable **CoQ redox pool**, which can buffer or limit throughput depending on regime (Burger et al., 2020). Third, cardiolipin-dependent **supercomplex organization**, treated as a state variable rather than a constant, motivated by cardiolipin’s documented role in supercomplex stability and formation (Zhang et al., 2002; Pfeiffer et al., 2003; Bazán et al., 2013). By unifying these elements, the syn-IMM framework aims to generate testable predictions about what dominates ATP output and stability, to define acceptance criteria for synthetic bioenergetic membranes, and to clarify which design knobs should be tuned first.

## Methodology

### Study design and objectives

This study develops a reproducible computational **synthetic inner mitochondrial membrane (syn-IMM)** framework to evaluate how coupled bioenergetic modules generate and dissipate **driving force**, and how those dynamics determine **ATP output**. The primary objective is **mechanistic interpretability**, meaning we deliberately isolate design variables that commonly covary in experimental reconstitution, including **membrane leak, lipid composition, organization state**, and **carrier pool constraints**, and quantify their individual and combined effects. The framework is positioned as a computational companion to experimental **de- and reconstruction** of mammalian respiratory function in defined membranes (Rimle et al., 2025).

### Model overview, two complementary simulators

To balance speed with mechanistic richness, we implemented two complementary dynamical simulators. **Model 1 (Δψ-proxy syn-IMM)** uses a single energetic state variable, **Δψ(t)**, as a proxy for membrane energization and computes **ATP synthesis** and **ATP export** directly from Δψ. This formulation is used for rapid perturbation tests, cardiolipin sweeps, Monte Carlo sensitivity analysis, and cardiolipin × leak landscapes. **Model 2 (multi-state syn-IMM)** explicitly tracks (i) **Δψ(t)** and **ΔpH(t)** as separable components of the **proton motive force (Δp)**, (ii) a finite **coenzyme Q (CoQ) redox pool**, and (iii) a cardiolipin-dependent **supercomplex assembly fraction S(t)** as a kinetic state. These additions address known interpretive pitfalls in bioenergetics where Δψ and ΔpH can decouple, carrier pools can saturate, and organization changes dynamically (Perry et al., 2011; Burger et al., 2020; Zhang et al., 2002).

### State variables, modules, and simulation workflow

In the multi-state model, the system is defined by four primary state variables, **Δψ(t)** (mV), **ΔpH(t)** (pH units), **q(t)** (the **QH2 fraction** of the total CoQ pool, bounded to [0, 1]), and **S(t)** (the **supercomplex fraction**, bounded to [0, 1]). The electrical-equivalent driving force is computed as **Δp(t) = Δψ(t) + 60·ΔpH(t)**, a standard conversion used for interpreting mitochondrial driving force and membrane potential probe behavior (Perry et al., 2011). This explicit partitioning allows the model to distinguish electrical versus chemical contributions to driving force, which is essential when interpreting perturbations or experimental conditions that alter buffering and ion gradients.

We represent syn-IMM function as a coupled set of modules that map to experimentally meaningful design levers. Electron flow is defined by a **redox bottleneck module** in which electron input reduces the CoQ pool and downstream oxidation consumes QH2. Two effective fluxes capture these processes, **J_red(t)** (CoQ reduction, upstream “Complex I-like” input) and **J_ox(t)** (CoQH2 oxidation, downstream “Complex III/IV-like” transfer). Net electron throughput is set by bottleneck logic, **J_e(t) = min(J_red(t), J_ox(t))**, ensuring that saturation on either side constrains throughput and naturally produces saturating redox regimes.

CoQ is modeled as a finite lipid-phase carrier pool with size **CoQ_total** (nmol CoQ per mg protein equivalent). The reduced fraction evolves according to **dq/dt = (J_red − J_ox)/CoQ_total**, which creates explicit saturation regimes, q(t) approaches 1 when oxidation is limiting and approaches 0 when reduction is limiting. This is consistent with the functional relevance and measurability of CoQ pool redox state in vivo and in reconstructed systems (Burger et al., 2020). Net throughput then drives **proton pumping**, represented as increases in both **Δψ** and **ΔpH** proportional to **J_e(t)**, scaled by a **pump gain** term. Pump gain is increased by the organization state **S(t)** and by a cardiolipin factor, capturing improved organization-dependent efficiency.

ATP synthesis is modeled as a steep nonlinear function of driving force, **J_ATP(t) = V_max · f(Δp)**, where f(Δp) is a sigmoid/Hill-type function that rises sharply near a midpoint. This reflects classic evidence that ATP synthesis is strongly dependent on transmembrane voltage and driving force and exhibits threshold-like behavior (Kaim & Dimroth, 1999). **Energetic back-action** is implemented by draining Δp in proportion to ATP synthesis, with the drain partitioned between **Δψ** and **ΔpH** using parameter **α**, so that Δψ and ΔpH can respond differently to coupling changes. This is necessary because Δψ-only interpretations can be misleading when ΔpH changes substantially under different buffering or conductance conditions (Perry et al., 2011).

Dissipation is represented by an explicit **leak module** with separate nonlinear leak functions for Δψ and ΔpH, implemented as negative terms in the Δψ and ΔpH dynamics. **Leak** is treated as a primary **engineering constraint** because it directly erodes Δp and therefore suppresses ATP output, consistent with general coupling logic used in mitochondrial assessment (Brand & Nicholls, 2011).

Export of ATP to an externally measurable pool is represented by a **transport gating module, J_export(t) = min(J_ATP(t), ANT_cap, PiC_cap)**, allowing explicit discrimination between **synthesis-limited** and **transport-limited** regimes. Functional reconstitution literature supports treating **ANT** as high-capacity in many liposome contexts unless deliberately restricted (Kreiter et al., 2020).

Cardiolipin effects are implemented in two coupled ways. First, a smooth cardiolipin **organization factor** scales effective organization-dependent throughput. Second, a reversible kinetic model treats organization as a dynamic state, **dS/dt = k_on(CL, q)·(1−S) − k_off(CL)·S**. This formulation is motivated by evidence that cardiolipin is required for supercomplex formation (Zhang et al., 2002), cardiolipin stabilizes respiratory chain supercomplexes (Pfeiffer et al., 2003), and cardiolipin-dependent reconstitution of supercomplexes can be demonstrated using purified complexes (Bazán et al., 2013). By making S(t) kinetic, the model produces explicit predictions about **recovery dynamics**, steady-state organization, and how organization interacts with leak and driving force.

Parameters were selected to satisfy three constraints. First, **qualitative wiring logic**, electron throughput increases Δp, Δp drives ATP synthesis, and leak diminishes Δp. Second, **plausible operating regimes**, Δp partitioning is computed using a standard conversion and interpreted in the context of practical guidance on potential probe use (Perry et al., 2011). Third, **biological grounding**, cardiolipin-supercomplex dependence and reconstitution feasibility are aligned with the cited supercomplex and reconstruction literature (Zhang et al., 2002; Pfeiffer et al., 2003; Bazán et al., 2013; Rimle et al., 2025). The intent is not exact quantitative prediction of any single preparation, but robust qualitative predictions with an interpretable sensitivity structure.

All simulations were implemented in Python using explicit time stepping (difference-equation integration) with fixed dt and multi-minute trajectories sufficient to approach quasi-steady state. Initial conditions represent an initially weakly energized membrane and a partially reduced carrier pool. Outputs are recorded as full time series and summarized at defined endpoints, enabling direct generation of tables.

To mimic experimental design logic, we performed a structured set of in silico experiments. A **baseline run** establishes a reference operating regime. **Component perturbations** modify one knob at a time, cardiolipin fraction, leak, ATP synthase capacity, transport caps, CoQ_total, and supercomplex kinetic rates, to isolate mechanistic signatures. A continuous **cardiolipin sweep** identifies a **performance window**, while **Monte Carlo sensitivity analysis** samples plausible parameter distributions to rank dominant drivers. Finally, **cardiolipin × leak landscapes** generate heatmaps of output to identify robust operating regions and collapse regimes.

All results are visualized directly in the notebooks using time courses, perturbation overlays, scatter plots, and heatmaps. Endpoint tables summarize scenario results and correlation tables rank sensitivity drivers. Model execution is deterministic for fixed parameters, Monte Carlo sampling is reproducible using fixed random seeds, and all figures and tables are regenerated end-to-end from notebook execution, ensuring complete reproducibility from code to manuscript outputs. Code availability is provided in GitHub repository (https://github.com/mpetalcorin/Synthetic-Inner-Mitochondrial-Membrane-syn-IMM-Simulator) under an open-source license.

## Results

### Baseline syn-IMM reaches an energized state and exports ATP

The baseline Δψ-proxy syn-IMM simulation shows progressive membrane energization and ATP production over 0-6 min. **Δψ rises** steadily (Figure 1A) and reaches **24.695 mV** at 6 min (Table 1). The **ATP synthesis rate** increases in parallel and reaches **0.6298 nmol·min**^**−1**^**·mg**^**−1**^ at 6 min (Figure 1B, Table 1). Under baseline transport settings, export is not limiting because the **export-limited rate overlaps the ATP synthase rate** (Figure 1B), resulting in accumulated exported ATP of **1.5840 nmol·mg**^**−1**^ by 6 min (Figure 1C, Table 1). These baseline outputs define the reference operating point for perturbation and landscape analyses.

**Table 1.** Baseline endpoint metrics (Δψ-proxy model, t = 6 min). Baseline steady-state values for **Δψ**, ATP synthesis/export rates, accumulated exported ATP, and cardiolipin factor, used as reference for all comparisons.

| Metric | Value |
| --- | --- |
| $\Delta\psi$ (mV) | 24.695 |
| ATP synthase rate (nmol·min<sup>-1</sup>·mg<sup>-1</sup>) | 0.6298 |
| Exported ATP rate (nmol·min<sup>-1</sup>·mg<sup>-1</sup>) | 0.6298 |
| Accumulated exported ATP (nmol·mg<sup>-1</sup>) | 1.5840 |
| Cardiolipin factor (unitless) | 1.000 |

**Figure 1.**
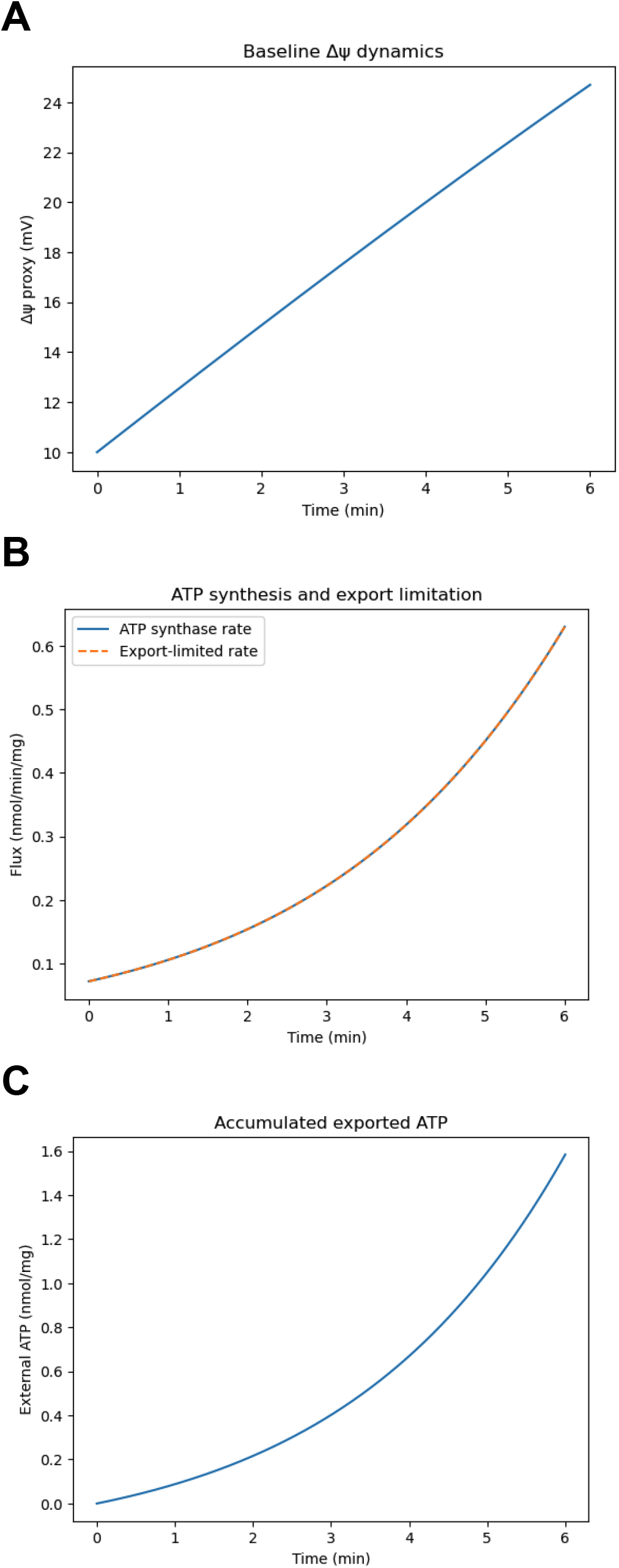
Baseline syn-IMM dynamics (Δψ-proxy model). Baseline time courses (0–6 min) showing (A) **Δψ** (mV), (B) **ATP synthesis rate** and **export-limited ATP rate** (nmol·min^−1^·mg^−1^), and (C) **accumulated exported ATP** (nmol·mg^−1^). Overlap of ATP synthesis and export-limited curves indicates export is not limiting under baseline settings.

### Component perturbations reveal distinct control points for energization and ATP throughput

Perturbation experiments demonstrate that different modules produce different signatures in **Δψ** and **ATP export** (Figure 2A-B; Table 2). Shifting cardiolipin away from the optimum reduces performance in a non-monotonic manner. **Low CL (5%)** reduces Δψ to **21.711 mV** and ATP export to **0.4088**, while **High CL (30%)** reduces Δψ to **22.093 mV** and ATP export to **0.4322** (Figure 2; Table 2), consistent with a **cardiolipin performance window** rather than linear scaling. Increasing dissipative loss (**High leak**) reduces ATP export to **0.5573** with a smaller reduction in Δψ to **23.847 mV** (Figure 2; Table 2), indicating leak primarily suppresses effective coupling and usable output. Reducing ATP synthase capacity yields the clearest “energized but unproductive” state, **Δψ remains near baseline** (24.950 mV) while **ATP export collapses** to **0.1815** (Figure 2; Table 2). Finally, **ANT limited** does not alter output in this parameterization, because ATP export remains synthesis-limited rather than transport-limited across the window (Figure 2B; Table 2).

**Table 2.** Perturbation endpoints (Δψ-proxy model, t = 6 min). Endpoint comparison across perturbations reporting **Δψ**, ATP synthesis rate, ATP export rate, accumulated exported ATP, and cardiolipin factor, quantifying distinct signatures for lipid shifts, leak increase, ATP synthase limitation, and transport limitation.

| Scenario | $\Delta\psi$ (mV) | ATP rate (nmol·min<sup>-1</sup>·mg<sup>-1</sup>) | ATP export (nmol·min<sup>-1</sup>·mg<sup>-1</sup>) | ATP_ext (nmol·mg<sup>-1</sup>) | CL factor |
| --- | --- | --- | --- | --- | --- |
| Baseline | 24.695 | 0.6298 | 0.6298 | 1.5840 | 1.000 |
| Low CL (5%) | 21.711 | 0.4088 | 0.4088 | 1.1846 | 0.799 |
| High CL (30%) | 22.093 | 0.4322 | 0.4322 | 1.2287 | 0.825 |
| High leak | 23.847 | 0.5573 | 0.5573 | 1.4749 | 1.000 |
| ANT limited | 24.695 | 0.6298 | 0.6298 | 1.5840 | 1.000 |
| Low ATP synthase | 24.950 | 0.1815 | 0.1815 | 0.4483 | 1.000 |

Units: ATP rate and ATP export in nmol·min<sup>-1</sup>·mg<sup>-1</sup>, ATP\_ext in nmol·mg<sup>-1</sup>

**Figure 2.**
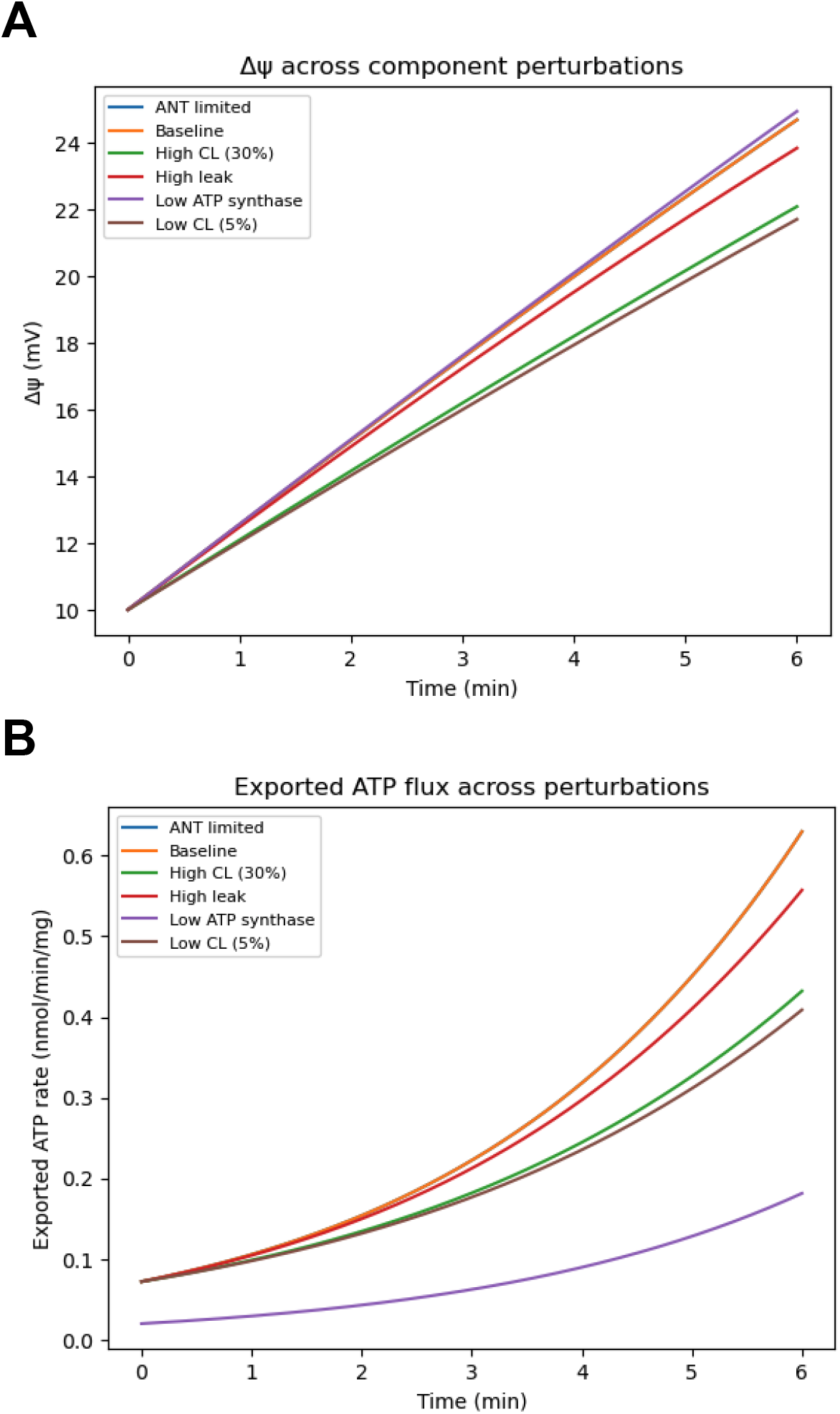
Component perturbations shift Δψ and ATP export (Δψ-proxy model). Overlay time courses (0-6 min) comparing **Baseline, Low CL (5%), High CL (30%), High leak, ANT limited**, and **Low ATP synthase** for (A) **Δψ** (mV) and (B) **exported ATP rate** (nmol min^−1^ mg^−1^), isolating distinct energization versus throughput failure modes.

### Cardiolipin sweep identifies the optimum that maximizes both Δψ and ATP export

A continuous cardiolipin sweep shows that both **exported ATP rate** and **Δψ** peak near the same cardiolipin fraction (Figure 3A-B). The maximum occurs at **CL = 0.180**, where **Δψ = 24.695 mV** and **ATP export = 0.6298 nmol·min**^**−1**^**·mg**^**−1**^ (Table 3). This coupling of maxima indicates that in the Δψ- proxy model, cardiolipin primarily acts by improving organization-dependent throughput that stabilizes energization, thereby increasing ATP output.

**Table 3.** Cardiolipin peak performance point (Δψ-proxy model). Cardiolipin fraction at which ATP export is maximal, with corresponding Δψ and ATP export values, summarizing the unimodal performance window.

| Quantity | Value |
| --- | --- |
| Cardiolipin fraction at peak (mol) | 0.180 |
| $\Delta\psi$ at peak (mV) | 24.695 |
| ATP export at peak (nmol·min<sup>-1</sup>·mg<sup>-1</sup>) | 0.6298 |

**Figure 3.**
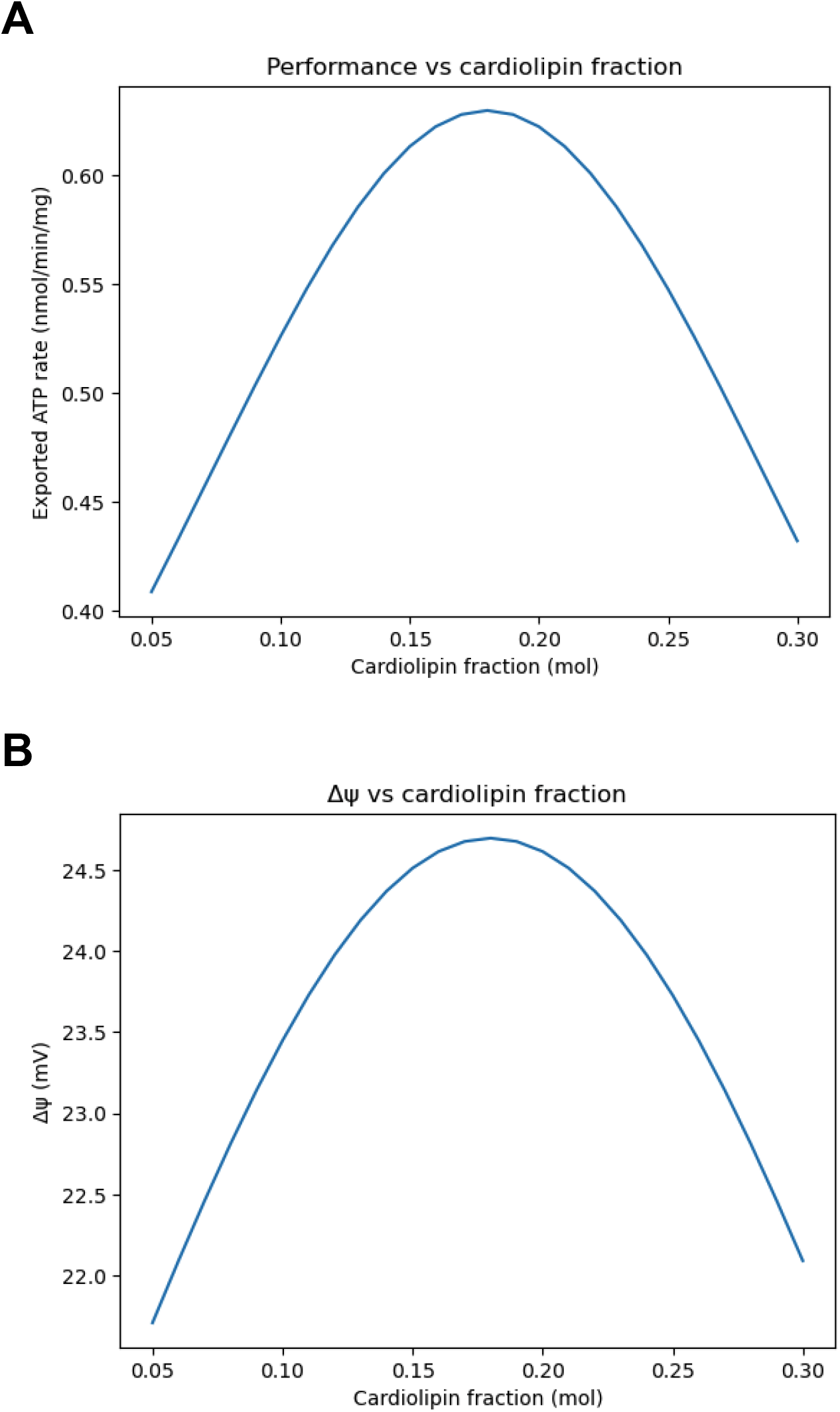
Cardiolipin titration reveals a performance window (Δψ-proxy model). Steady-state (t = 6 min) outputs plotted versus **cardiolipin fraction** (0.05–0.30 mol) for (A) **exported ATP rate** (nmol·min^−1^·mg^−1^) and (B) **Δψ** (mV), showing a unimodal optimum near CL ≈ 0.18.

### Monte Carlo sampling ranks dominant drivers of ATP output and shows cardiolipin is non-monotonic

Monte Carlo sampling reveals broad dispersion in ATP output (Figure 4), with clear suppression as leak increases (Figure 4A) and increased output with higher ATP synthase capacity (Figure 4B). The scatter against raw cardiolipin fraction is weak (Figure 4C) because cardiolipin acts through a peaked window rather than monotonic scaling. Correlation ranking confirms that ATP export aligns most strongly with Δψ and respiratory capacity (J_e_max), followed by ATP synthase capacity, while leak and energetic drain terms contribute negatively (Table 4).

**Table 4.** Monte Carlo correlation ranking for ATP export (Δψ-proxy model, n = 2000). Pearson correlations between sampled parameters/derived variables and ATP export rate, providing a practical dominance ranking of drivers of output variance.

| Rank | Variable | r with ATP export |
| --- | --- | --- |
| 1 | $\Delta\psi$ (mV) | 0.8187 |
| 2 | J_e_max (respiratory capacity) | 0.8078 |
| 3 | ATP_Vmax (ATP synthase capacity) | 0.3847 |
| 4 | CL factor (unitless) | 0.1496 |
| 5 | leak_g (unitless) | -0.0860 |
| 6 | mV_per_ATP (unitless) | -0.0370 |
| 7 | PiC_cap (nmol·min<sup>-1</sup>·mg<sup>-1</sup>) | 0.0249 |
| 8 | ANT_cap (nmol·min<sup>-1</sup>·mg<sup>-1</sup>) | 0.0171 |
| 9 | cardiolipin fraction (mol) | -0.0069 |

**Figure 4.**
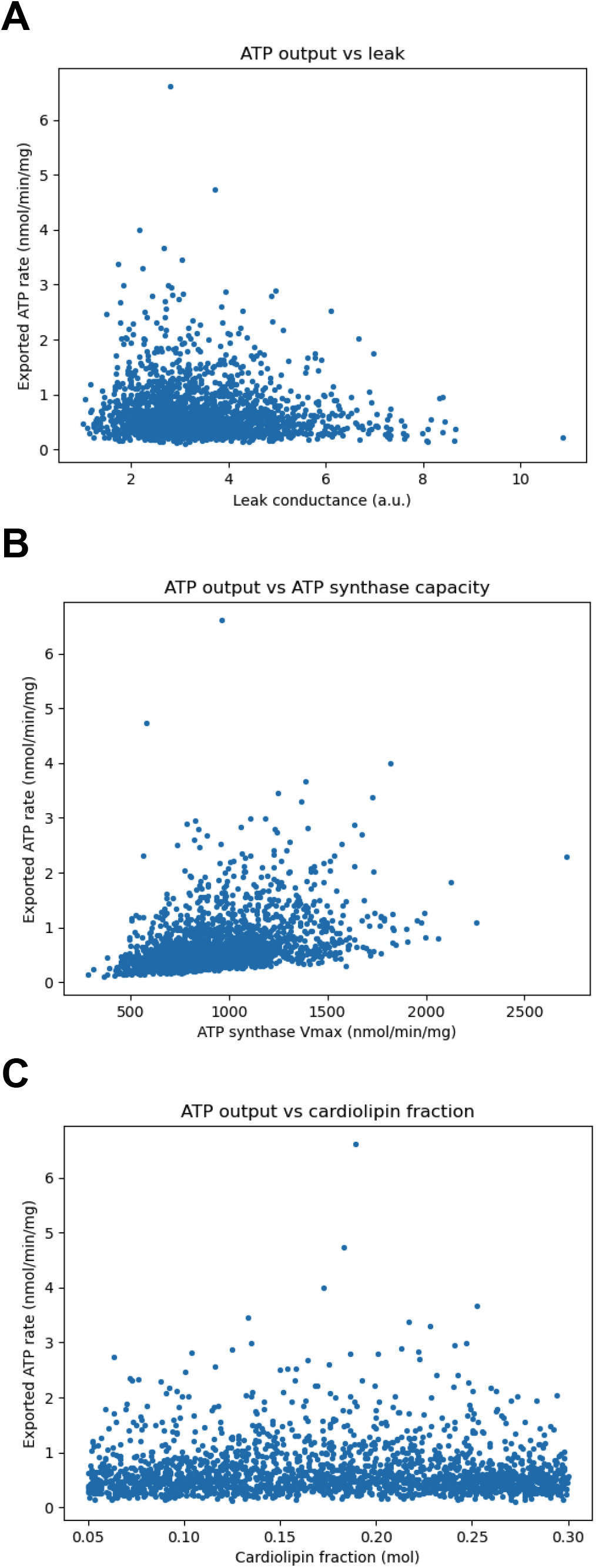
Monte Carlo sensitivity of ATP output (Δψ-proxy model). Scatter plots from Monte Carlo sampling (n = 2000) showing exported ATP rate versus (A) **leak conductance**, (B) **ATP synthase capacity (Vmax)**, and (C) **cardiolipin fraction**, visualizing dominant drivers and the non-monotonic lipid dependence.

### Cardiolipin × leak landscapes define an operating band that collapses under uncoupling

Two-factor heatmaps generalize the perturbation results. A high-performance band centered near cardiolipin ∼0.18 is visible for both ATP export and Δψ (Figure 5A-B). This band is progressively suppressed as leak increases, demonstrating that cardiolipin optimization enhances performance primarily within coupling-permissive regimes, while high leak globally constrains Δψ stability and ATP output.

**Figure 5.**
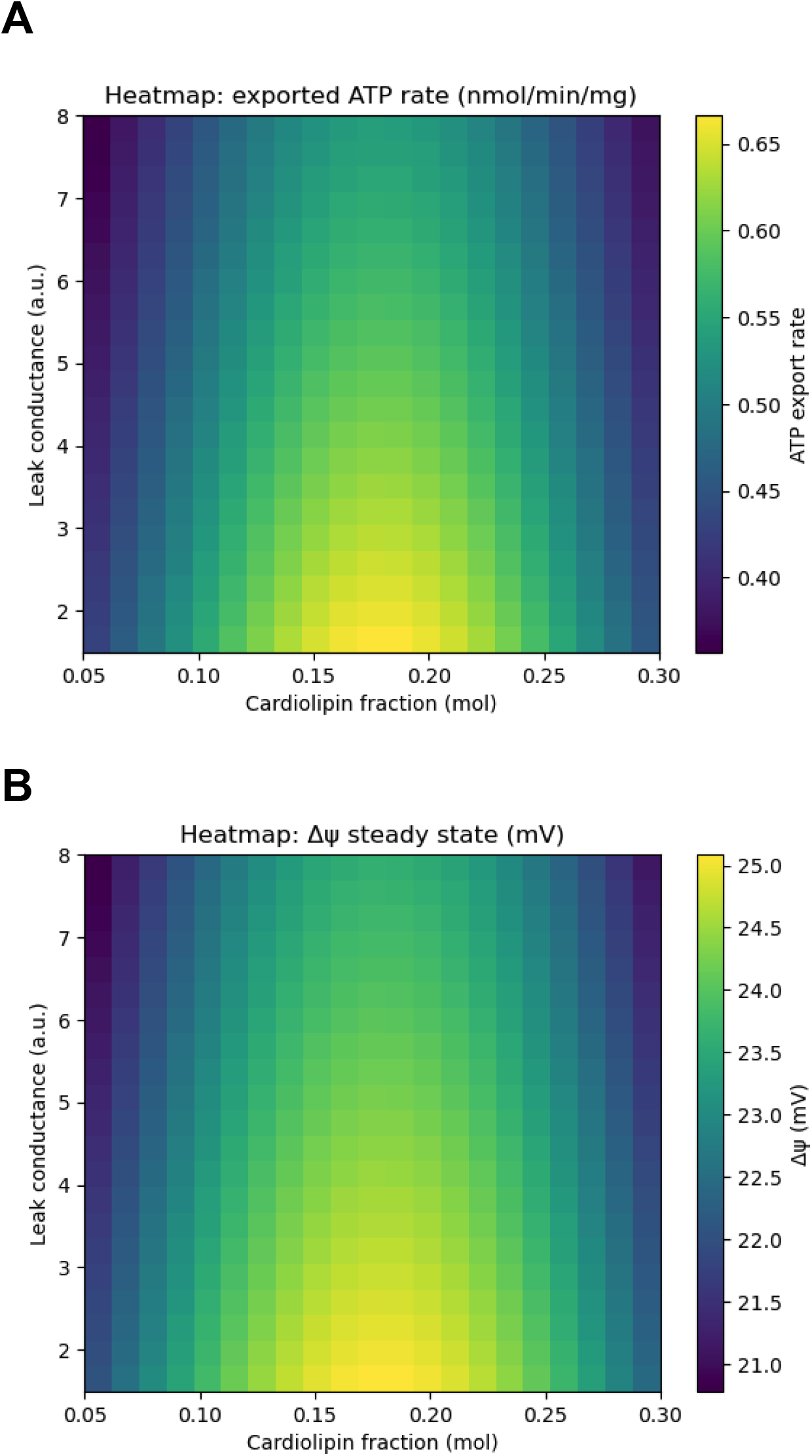
Two-factor landscape, cardiolipin × leak tradeoff (Δψ-proxy model). Heatmaps over **cardiolipin fraction** (x-axis) and **leak conductance** (y-axis) showing steady-state (t = 6 min) for (A) **exported ATP rate** (nmol·min^−1^·mg^−1^) and (B) **Δψ** (mV), identifying a high-performance band near CL ≈ 0.18 suppressed by increasing leak.

### Δp partitioning shows distinct Δψ and ΔpH dynamics rather than a single “potential”

The multi-state model explicitly tracks **Δψ, ΔpH**, and **Δp = Δψ + 60·ΔpH** (Figure 6). In the baseline, Δp is dominated by Δψ, while 60·ΔpH contributes a smaller additive component. The baseline time-series table confirms this partition evolves over time (Table 6), enabling mechanistic discrimination between electrical versus chemical contributions to Δp, which is not possible in the single-variable Δψ proxy.

**Figure 6.**
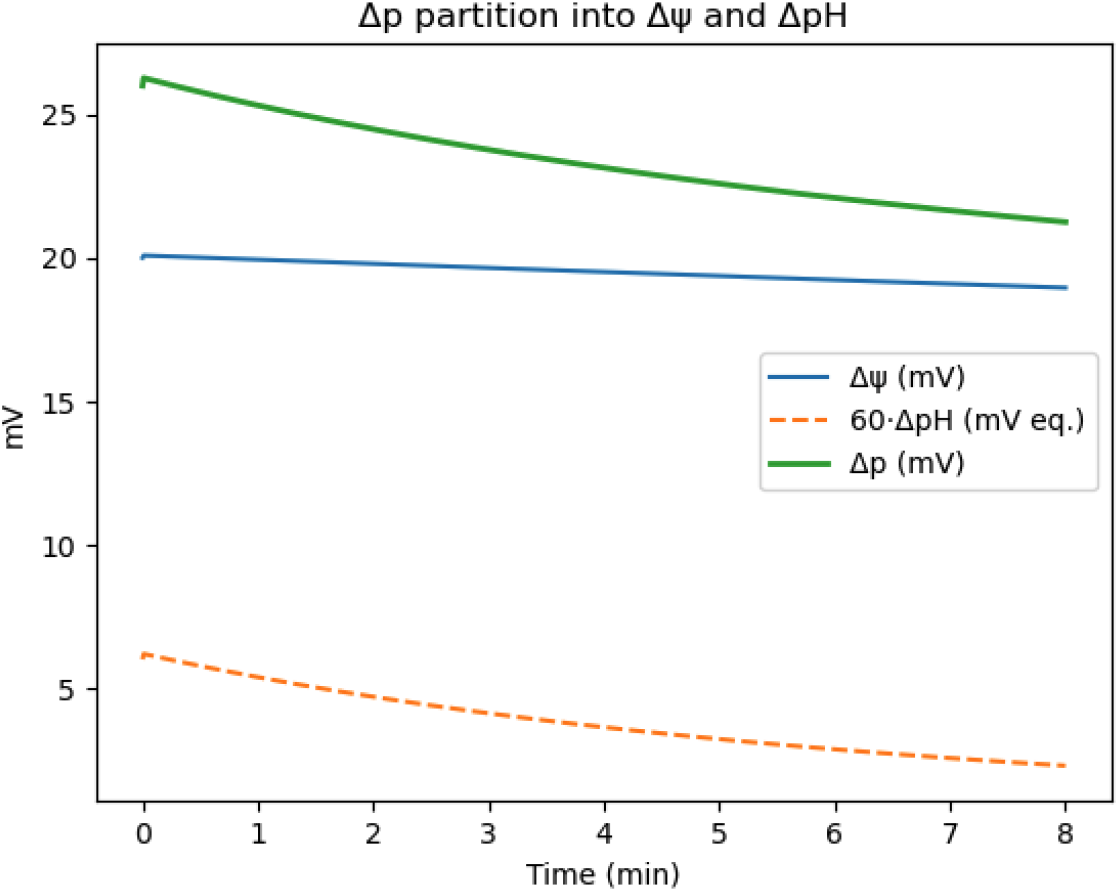
Δp partitioning into Δψ and ΔpH (multi-state model). Baseline time courses (0-8 min) of **Δψ** (mV), **60·ΔpH** (mV equivalent), and **Δp** (mV), computed as **Δp = Δψ + 60·ΔpH**, showing distinct evolution of electrical and chemical components.

### Supercomplex fraction S(t) rises kinetically, but ATP export does not necessarily increase with S

In the multi-state baseline, **S(t)** increases from ∼0.30 toward ∼0.80 (Figure 7A; Table 6), demonstrating organization is a **kinetic state** rather than a fixed property. ATP export shows its own trajectory (Figure 7B), supporting the system-level point that increasing organization does not guarantee increasing ATP flux if the dominant constraint is Δp erosion or other dissipative terms.

**Figure 7.**
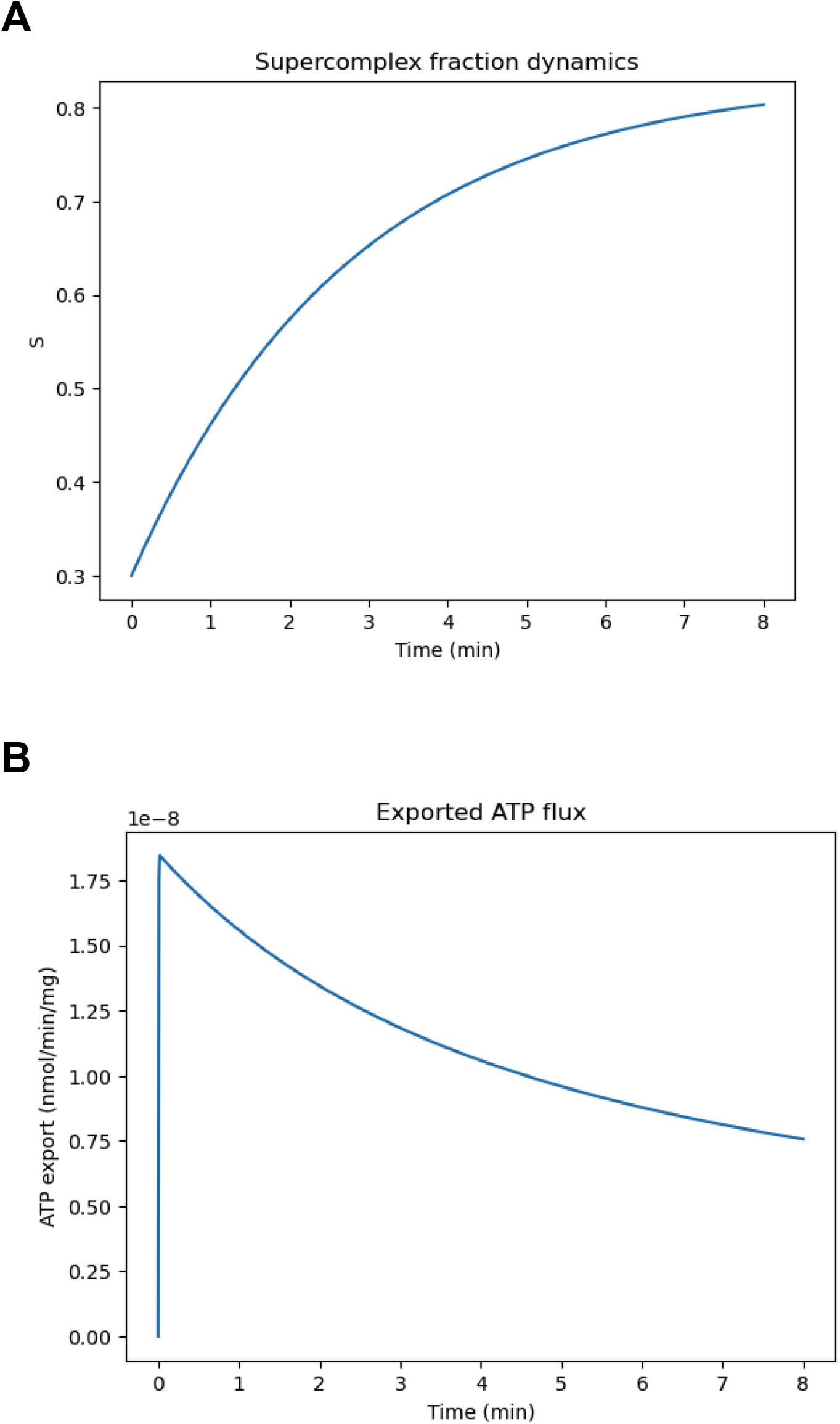
Supercomplex assembly kinetics and ATP export (multi-state baseline). Baseline time courses (0–8 min) showing (A) **supercomplex fraction S(t)** and (B) **exported ATP flux** (nmol·min^−1^·mg^−1^), linking time-dependent organization to functional output.

### Multi-state perturbations separate leak-dominant defects from organization-dominant and assembly-kinetic effects

The perturbation panel demonstrates that **High leak** produces the strongest suppression of Δp and ATP export while leaving S similar to baseline in this run (Figure 8A-C; Table 5), confirming that leak remains the dominant antagonist even when organization is high. Cardiolipin changes strongly shift S, with S reduced to **0.545** at Low CL and **0.576** at High CL (Table 5), showing lipid-dependent organization sensitivity. Altered supercomplex kinetics change S substantially, **0.483** for weak assembly and **0.948** for strong assembly (Table 5), providing a direct organizational lever, even though the ATP export differences are comparatively small under these particular parameter settings.

**Table 5.** Multi-state perturbation endpoints (t = 8 min). End-of-run values for **Δψ, ΔpH, Δp, QH2 fraction, supercomplex fraction S**, ATP export flux, and accumulated exported ATP across multi-state perturbations.

| Scenario | $\Delta\psi$ (mV) | $\Delta\text{pH}$ | $\Delta\text{p}$ (mV) | QH2 fraction | S (supercomplex) | ATP export (nmol·min<sup>-1</sup>·mg<sup>-1</sup>) | ATP_ext (nmol·mg<sup>-1</sup>) |
| --- | --- | --- | --- | --- | --- | --- | --- |
| Baseline | 18.956 | 0.0382 | 21.247 | 0.000 | 0.804 | 7.5628E-09 | 9.0704E-08 |
| Low CL (5%) | 18.940 | 0.0380 | 21.219 | 0.000 | 0.545 | 7.5255E-09 | 9.0028E-08 |
| High CL (30%) | 18.941 | 0.0380 | 21.222 | 0.000 | 0.576 | 7.5293E-09 | 9.0095E-08 |
| High leak | 17.596 | 0.0153 | 18.514 | 0.000 | 0.804 | 4.6533E-09 | 6.6204E-08 |
| Low CoQ pool | 18.956 | 0.0382 | 21.247 | 0.000 | 0.804 | 7.5628E-09 | 9.0704E-08 |
| High CoQ pool | 18.956 | 0.0382 | 21.247 | 0.000 | 0.804 | 7.5628E-09 | 9.0704E-08 |
| Weak supercomplex assembly | 18.956 | 0.0382 | 21.247 | 0.000 | 0.483 | 7.5626E-09 | 9.0701E-08 |
| Strong supercomplex assembly | 18.956 | 0.0382 | 21.247 | 0.000 | 0.948 | 7.5630E-09 | 9.0708E-08 |

**Table 6.** Multi-state baseline trajectory, representative start and end points (t = 0 and 8 min). Baseline initial and final states reporting **Δψ, ΔpH, Δp, QH2 fraction, S**, redox flux proxies, ATP synthesis/export flux, and accumulated ATP, illustrating trajectory evolution and endpoint consistency.

| t (min) | $\Delta\psi$ (mV) | $\Delta\text{pH}$ | $\Delta\text{p}$ (mV) | QH2 fraction | S | J_red (nmol·min<sup>-1</sup>·mg<sup>-1</sup>) | J_ox (nmol·min<sup>-1</sup>·mg<sup>-1</sup>) | J_e (nmol·min<sup>-1</sup>·mg<sup>-1</sup>) | J_ATP (nmol·min<sup>-1</sup>·mg<sup>-1</sup>) | J_export (nmol·min<sup>-1</sup>·mg<sup>-1</sup>) | ATP_ext (nmol·mg<sup>-1</sup>) |
| --- | --- | --- | --- | --- | --- | --- | --- | --- | --- | --- | --- |
| 0.00 | 20.000 | 0.1000 | 26.000 | 0.300 | 0.300 | 0.0000E+00 | 0.0000E+00 | 0.0000E+00 | 0.0000E+00 | 0.0000E+00 | 0.0000E+00 |
| 8.00 | 18.956 | 0.0382 | 21.247 | 0.000 | 0.804 | 0.0000E+00 | 1.1325E+03 | 0.0000E+00 | 7.5628E-09 | 7.5628E-09 | 9.0704E-08 |

**Figure 8.**
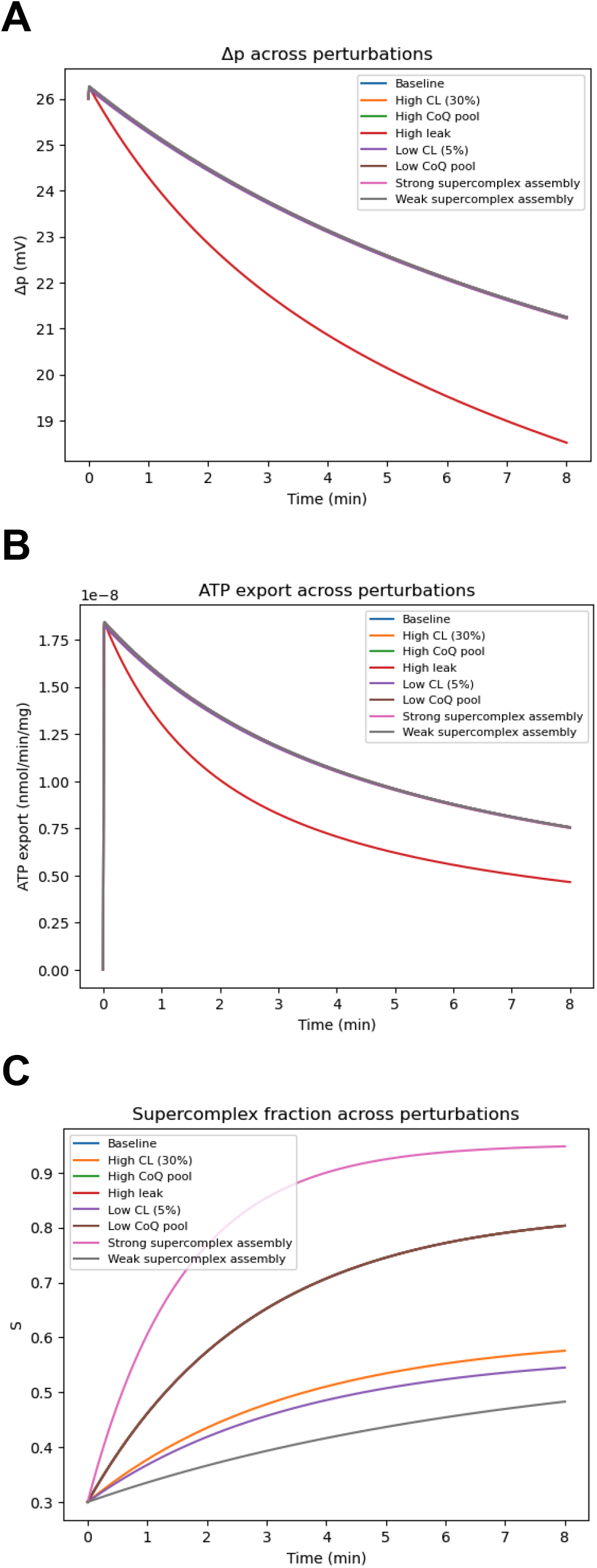
Multi-state perturbation panel, Δp, ATP export, and S(t). Overlay plots (0–8 min) comparing **Baseline, Low CL (5%), High CL (30%), High leak, Low CoQ pool, High CoQ pool, Weak supercomplex assembly**, and **Strong supercomplex assembly** for (A) **Δp** (mV), (B) **ATP export** (nmol·min^−1^·mg^−1^), and (C) **S(t)**, distinguishing driving-force, carrier, and organization effects.

### Multi-state cardiolipin × leak landscapes show organization has a lipid window but remains constrained by leak

The multi-state heatmaps extend these results into a global landscape. ATP export decreases with leak across cardiolipin values (Figure 9A), consistent with Δp erosion (Figure 9B). The supercomplex fraction S exhibits a cardiolipin-centered band where organization is highest (Figure 9C), confirming a lipid-defined organization window. However, comparing panels shows that even high S does not guarantee high ATP export at high leak, reinforcing the hierarchy established earlier, **coupling integrity is required for organization-dependent gains**.

**Figure 9.**
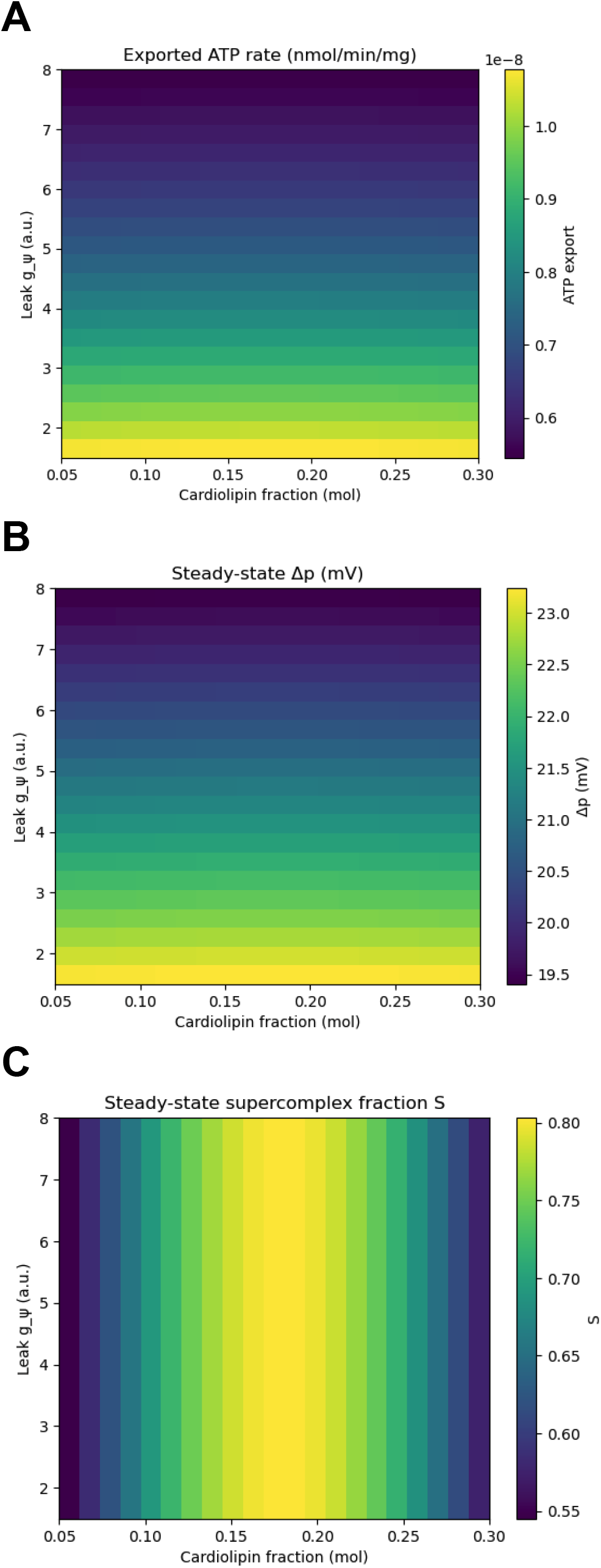
Cardiolipin × leak landscapes (multi-state model). Heatmaps over **cardiolipin fraction** (x-axis) and **leak gψ** (y-axis) showing steady-state (t = 8 min) for (A) **ATP export** (nmol·min^−1^·mg^−1^), (B) **Δp** (mV), and (C) **supercomplex fraction S**, demonstrating that a lipid-defined organization window remains constrained by leak.

## 4. Discussion

Experimental studies have shown that key parts of mammalian **oxidative phosphorylation** can be taken apart and rebuilt in defined membranes, which means it is now realistic to ask what a “synthetic” energy membrane needs in order to make ATP (Rimle et al., 2025). Because many experimental variables change together in real reconstitutions, such as how leaky the membrane is, which lipids are present, how well the complexes organize, and how much carrier is available, a computational syn-IMM is useful as a clean test bed. The main value of this model is not to claim perfect absolute rates, but to help interpret patterns, for example, what it means when **Δψ, Δp**, ATP flux, and organization change in a particular combination, and which constraint is most likely limiting.

A common mistake in bioenergetics is to assume that a high **membrane potential (Δψ)** automatically means high ATP production. In reality, ATP output depends on whether the system is well coupled and whether downstream machinery has enough capacity to convert the driving force into ATP (Brand & Nicholls, 2011). In our simulations, we see a clear “energized but unproductive” situation, Δψ can remain relatively high while ATP export drops sharply when **ATP synthase capacity** is limiting. This matters for experimental troubleshooting, if Δψ looks acceptable but ATP output is low, the simplest explanations are often insufficient ATP synthase capacity, substrate gating, or transport limits, rather than an immediate failure of electron transfer.

Across perturbations, sensitivity sampling, and cardiolipin × leak maps, **leak** emerges as the strongest overall suppressor of ATP output because it dissipates the electrochemical driving force that ATP synthase requires. This is consistent with established coupling principles, when proton conductance increases or the membrane becomes partially uncoupled, ATP production falls even if electron transport can still proceed (Brand & Nicholls, 2011). The practical consequence is that membrane “tightness” is a hard engineering requirement for any syn-IMM, fine-tuning lipid composition or organization will have limited benefit if leak is not controlled first.

The lipid **cardiolipin** produces a clear “performance window” in the model, with an optimum region where both energization and ATP export are maximized. This is biologically plausible because cardiolipin is tightly linked to respiratory chain architecture. Classic work shows cardiolipin is required for robust **supercomplex** formation (Zhang et al., 2002), cardiolipin stabilizes respiratory supercomplexes (Pfeiffer et al., 2003), and purified-complex reconstitution can reproduce cardiolipin-dependent stabilization of higher-order assemblies (Bazán et al., 2013). In design terms, cardiolipin behaves like a tunable control knob that improves organization and stability, but only within a certain range, and only when leak is not dominating.

A key improvement in the multi-state extension is treating organization as a dynamic process rather than a static label. The supercomplex fraction **S(t)** rises over time and shifts under cardiolipin and assembly-kinetic perturbations, which mirrors the fact that many experimental questions are kinetic, how quickly organization forms, how stable it is, and how it behaves under stress. The model also predicts an important separation, supercomplex organization can look high while ATP output remains low if leak is high. This means supercomplex readouts alone cannot guarantee productive coupling, organization helps, but it cannot replace an intact driving force in a leaky system (Brand & Nicholls, 2011; Zhang et al., 2002; Pfeiffer et al., 2003).

Separating **Δp** into **Δψ** and **ΔpH** is also important because those two components can change differently. Practical guidance on mitochondrial potential probes emphasizes that Δψ-based measurements can be misleading if the pH gradient component shifts, because Δψ and ΔpH respond differently depending on buffering and ion fluxes (Perry et al., 2011). By tracking **Δp = Δψ + 60·ΔpH**, the model can distinguish whether loss of driving force is mainly due to electrical collapse, chemical gradient collapse, or both, which is especially useful in reconstituted systems where buffer conditions and ionic composition are controlled variables.

The explicit **CoQ pool** in the multi-state model highlights why a finite carrier pool often affects **robustness** and transient behavior more than the final steady plateau. CoQ pool size and redox state are measurable and biologically meaningful, supported by sensitive assays of CoQ redox state in vivo (Burger et al., 2020). In the parameter regime used for your current runs, changing CoQ pool size does not shift endpoints, which is itself informative, it implies that other constraints, such as leak and organization, dominate in that regime. In regimes closer to redox bottlenecks, however, a finite CoQ pool can saturate and create transient limitations, and the model is designed to reveal those cases.

The mitochondrial genetic architecture also explains why the model treats assembled complexes as the starting point. The human mitochondrial genome encodes key membrane-embedded OXPHOS subunits that form the hydrophobic core of the machinery (Anderson et al., 1981), which helps explain why many syn-IMM strategies begin with purified assembled complexes rather than attempting to express and insert those hydrophobic cores de novo. As synthetic biology moves toward expression-coupled membrane systems, future models will need to add explicit assembly constraints and insertion bottlenecks, and this framework provides a scaffold for that.

Finally, membrane geometry is a natural next frontier. The IMM is not only a biochemical surface, it is a shaped topology, and ATP synthase dimers can form rows that induce membrane curvature, linking organization directly to structure (Blum et al., 2019). Although our current models do not explicitly include curvature or diffusion constraints, the results suggest when geometry will matter most, after leak is controlled and organization is high, diffusion limitations and microdomain effects are likely to become the next-order constraints. Extending the syn-IMM model to include geometry would therefore be a logical step.

Overall, the figures and tables support a practical design order for synthetic bioenergetic membranes. First, minimize **leak** because it determines whether Δp can be maintained (Brand & Nicholls, 2011). Second, match **ATP synthase capacity** to pumping to avoid “energized but unproductive” states (Kaim & Dimroth, 1999). Third, tune **cardiolipin** to the window that stabilizes organization and improves throughput (Zhang et al., 2002; Pfeiffer et al., 2003; Bazán et al., 2013). Fourth, use **S(t)** kinetics and **Δp partitioning** to interpret time dependence and separate electrical from chemical limitations (Perry et al., 2011). Finally, once these fundamentals are satisfied, **carrier pools** and **geometry** can become dominant in specific regimes, motivating the next generation of syn-IMM models and experiments (Burger et al., 2020; Blum et al., 2019).

## 5. Conclusion

This study introduces a reproducible computational **synthetic inner mitochondrial membrane (syn-IMM)** framework that treats energy conversion as a coupled system, rather than a simple parts list.

Across baseline simulations, targeted perturbations, parameter sweeps, and sensitivity analyses, the model shows that ATP output depends on whether the membrane can maintain a stable driving force and whether downstream catalytic capacity can convert that driving force into ATP. The most consistent result is that **membrane leak** is the dominant constraint, because increased proton conductance dissipates the energetic gradient and compresses the feasible operating space for ATP production. Once leak is controlled, **ATP synthase capacity** becomes the main ceiling on ATP flux, explaining the “energized but low-ATP” regime in which membrane potential can remain relatively high while ATP output is suppressed when the ATP-producing module is limiting. The simulations also identify a cardiolipin-dependent **performance window**, consistent with cardiolipin’s role in stabilizing respiratory organization, where energization and ATP export are jointly optimized under coupling-permissive conditions. The multi-state extension strengthens mechanistic interpretation by explicitly partitioning driving force into **Δψ and ΔpH**, incorporating a finite **CoQ redox pool**, and representing organization as a dynamic **supercomplex state S(t)**, enabling clearer diagnosis of whether limitations arise from dissipation, catalytic capacity, organization kinetics, or carrier-pool bottlenecks. Overall, the syn-IMM platform provides a quantitative and interpretable bridge between reconstituted membrane experiments and system-level design, offering practical acceptance criteria and design priorities for engineering programmable bioenergetic membranes.

